# GR13-type plasmids in *Acinetobacter* potentiate the accumulation and horizontal transfer of diverse accessory genes

**DOI:** 10.1101/2022.01.12.475240

**Authors:** Robert A. Moran, Haiyang Liu, Emma L. Doughty, Xiaoting Hua, Elizabeth A. Cummins, Tomas Liveikis, Alan McNally, Zhihui Zhou, Willem van Schaik, Yunsong Yu

## Abstract

Carbapenem resistance and other antibiotic resistance genes (ARGs) can be found in plasmids in *Acinetobacter*, but many plasmid types in this genus have not been well-characterised. Here we describe the distribution, diversity and evolutionary capacity of *rep* group 13 (GR13) plasmids that are found in *Acinetobacter* species from diverse environments. Our investigation was prompted by the discovery of two GR13 plasmids in *A. baumannii* isolated in an intensive care unit (ICU). The plasmids harbour distinct accessory genes: pDETAB5 contains *bla*_NDM-1_ and genes that confer resistance to four further antibiotic classes, while pDETAB13 carries putative alcohol tolerance determinants. Both plasmids contain multiple *dif* modules, which are flanked by p*dif* sites recognised by XerC/XerD tyrosine recombinases. The ARG-containing *dif* modules in pDETAB5 are almost identical to those found in pDETAB2, a GR34 plasmid from an unrelated *A. baumannii* isolated in the same ICU a month prior. Examination of a further 41 complete, publicly available plasmid sequences revealed that the GR13 pangenome consists of just four core but 1086 accessory genes, 123 in the shell and 1063 in the cloud, reflecting substantial capacity for diversification. The GR13 core genome includes genes for replication and partitioning, and for a putative tyrosine recombinase. Accessory segments encode proteins with diverse putative functions, including for metabolism, antibiotic/heavy metal/alcohol tolerance, restriction-modification, an anti-phage system and multiple toxin-antitoxin systems. The movement of *dif* modules and actions of insertion sequences play an important role in generating diversity in GR13 plasmids. Discrete GR13 plasmid lineages are internationally disseminated and found in multiple *Acinetobacter* species, which suggests they are important platforms for the accumulation, horizontal transmission and persistence of accessory genes in this genus.

**Impact statement:** *Acinetobacter* species are particularly well-adapted for persistence in hospital environments where they pose a life-threatening infection risk to the most clinically-vulnerable patients. Plasmids with the potential to transfer multiple antibiotic resistance determinants between *Acinetobacter* species are therefore concerning, but most are not well-characterised. This work sheds further light on the poorly-understood mobile gene pool associated with *Acinetobacter*. We show here that GR13 plasmids carry a small set of core genes but have access to a highly diverse set of accessory segments that might provide fitness advantages under certain conditions. The complex evolutionary dynamics of GR13 plasmids appear to be driven by the exchange of *dif* modules and by the actions of a diverse population of insertion sequences. The novel *dif* modules characterised here emphasise the broader importance of these elements to the dissemination of accessory genes in *Acinetobacter*. This study has improved our understanding of the diversity and distribution of *dif* modules, plasmids that carry them, and how both disseminate in the continuum of *Acinetobacter* populations that link hospitals and the wider environment.

## Introduction

*Acinetobacter* is a genus of Gram-negative coccobacilli that are typically found in soils and moist environments but are also well-adapted for persistence in hospital settings (Visca *et al*., 2011). *A. baumannii* is the most prominent pathogenic species and can cause human infections with high mortality rates, particularly given some strains exhibit extensive antibiotic resistance that severely compromises treatment (Hamidian and Nigro, 2019; Visca *et al*., 2011). While less commonly reported, drug-resistant infections caused by other *Acinetobacter* species are an emerging threat (Endo *et al*., 2014; Mittal *et al*., 2015; Sieswerda *et al*., 2017; Silva *et al*., 2018; Yang *et al*., 2021).

Plasmids in *Acinetobacter* are typed according to replication initiation gene (*rep*) identity (Bertini *et al*., 2010). A recent review listed 33 *rep* groups (GRs) (Salgado-Camargo *et al*., 2020), and we have since described an additional group, GR34 (Liu *et al*., 2021). Plasmids carrying clinically-significant antibiotic resistance genes (ARGs) have been reported in *A. baumannii* (Blackwell and Hall, 2017; Hamidian *et al*., 2016; Hamidian and Hall, 2014; Liu *et al*., 2015; Nigro and Hall, 2014) and in other *Acinetobacter* species (Alattraqchi *et al*., 2021; Hayashi *et al*., 2021; Li *et al*., 2021; Silva *et al*., 2018; Yang *et al*., 2021), clearly indicating their important role in the emergence and transmission of antimicrobial resistance in this genus. Few plasmid groups have been the subject of comparative analyses, so how the remaining types evolve or are distributed, geographically and throughout the *Acinetobacter* genus, is poorly understood and their genetic structures remain largely undescribed.

Some *Acinetobacter* plasmids carry multiple pairs of recombination sites that resemble chromosomal *dif* sites, which are targets for XerC and XerD tyrosine recombinases (Balalovski and Grainge, 2020). These have been called plasmid-*dif* (p*dif*) sites (Blackwell and Hall, 2017), and have been shown to be recognised by *A. baumannii* XerC and XerD (Lin *et al*., 2020). ARGs have been found in p*dif*-flanked structures called *dif* modules, including the carbapenemase genes *bla*_OXA-24_ (D’Andrea *et al*., 2009), *bla*_OXA-58_ (Bertini *et al*., 2007), and *bla*_OXA-72_ (Kuo *et al*., 2016), the tetracycline resistance gene *tet*(39) (Blackwell and Hall, 2017), the macrolide resistance genes *msr*(E)-*mph*(E) (Blackwell and Hall, 2017), the aminoglycoside resistance gene *aacC2d* and the sulphonamide resistance gene *sul2* (Liu *et al*., 2021). Identical ARG-containing *dif* modules have been found in multiple contexts and in different types of plasmids (Blackwell and Hall, 2017). Further *dif* modules, including those carrying genes for chromium resistance, a serine recombinase, RND efflux system and multiple toxin-antitoxin systems have also been described (Blackwell and Hall, 2017; Hamidian and Hall, 2018; Liu *et al*., 2021; Mindlin *et al*., 2018). Given the apparent importance of *dif* modules to the evolution of some *Acinetobacter* plasmids, it is important to understand the breadth of genetic cargo they carry and which types of plasmids can interact with them.

We recently described the GR34 family of plasmids that share a 10 kbp core segment but can grow to as large as 190 kbp through the acquisition of *dif* modules (Liu *et al*., 2021). The exemplar GR34 plasmid, pDETAB2, is from an *A. baumannii* isolated in an intensive care unit (ICU) in Hangzhou, China, and carries six ARGs in a series of *dif* modules (Liu *et al*., 2021). Here, we report two GR13-type plasmids found in unrelated *A. baumannii* isolated one and two months later in that same ICU, one cryptic and the other carrying ARG-containing *dif* modules identical to ones in pDETAB2. In order to contextualise the differences between them, we undertook a detailed comparative analysis of the ICU GR13 plasmids and 41 complete GR13 plasmid sequences from GenBank. This facilitated the first evaluation of the distribution, gene content, structures and evolutionary characteristics of the GR13 plasmid family.

## Materials and Methods

### Ethics

Ethical approval and informed consent were obtained by the Sir Run Run Shaw Hospital local ethics committee (approval number 20190802-1).

### Bacterial isolation and antibiotic susceptibility testing

DETAB-E227 was isolated from a cleaning cart surface swab and DETAB-P39 from a patient rectal swab in Sir Run Run Shaw Hospital Intensive Care Unit in Hangzhou, China in 2019. Both samples were cultured on CHROMagar (CHROMagar, Paris, France) containing 2 mg/L meropenem at 37°C for 24 hours. Isolated colonies of presumptive *A. baumannii* were sub-cultured on Mueller-Hinton agar (MHA) (Oxoid, Hampshire, UK) and incubated at 37°C for 24 hours. MICs for imipenem, meropenem, tobramycin, gentamicin, ciprofloxacin, levofloxacin, ceftazidime, colistin and tigecycline were determined using broth microdilution with results interpreted according to CLSI 2019 guidelines.

### Plasmid transfer assays

DETAB-E227 was filter-mated with a rifampicin-resistant derivative of *A. baumannii* ATCC 17978 or *A. nosocomialis* strain XH1816 as described previously (Jin *et al*., 2018). XH1816 is a colistin-resistant, meropenem-sensitive clinical *A. nosocomialis* strain XH1816 that was isolated from a human urine sample in 2011. Transconjugants were selected on MHA supplemented with rifampicin (50 µg/mL) and meropenem (8 µg/mL). The identity of transconjugants was confirmed through PFGE fingerprinting after digestion of genomic DNA with *ApaI*. Transconjugants were tested for the presence of pDETAB4 and pDETAB5 by PCR with primers that target the replication genes of each plasmid (Table S1). ATCC 17978 transconjugants containing pDETAB2 were mated with XH1816 as above, with transconjugants selected on MHA supplemented with colistin (2 µg/mL) and meropenem (8 µg/mL).

### S1 nuclease digestion, pulsed field gel electrophoresis and Southern blot

To confirm transfer had occurred, plasmids were visualised following S1 nuclease treatment via PFGE, and the locations of resistance genes were confirmed via Southern blot as described previously (Quan *et al*., 2017). Briefly, genomic DNA was digested with S1 nuclease (TaKaRa, Kusatsu, Japan) at 37°C for 20 minutes. Treated DNA was loaded on a 1% agarose Gold gel and PFGE was performed at 14°C for 18 hours, with 6 V/cm and pulse times from 2.16 to 63.8 seconds using the Bio-Rad CHEF-Mapper XA machine (Bio-Rad, California, USA). DNA was transferred to a positively-charged nylon membrane (Millipore, Billerica, MA, USA) by the capillary method and hybridised with digoxigenin-labelled *bla*_OXA- 58_ and *bla*_NDM-1_-specific probes with an NBT/BCIP colour detection kit (Roche, Mannheim, Germany) according to the manufacturer’s instructions. *Xba*I-treated genomic DNA from *Salmonella enterica* H9812 was used as a size marker.

### Whole genome sequencing and analysis

Genomic DNA was extracted from *A. baumannii* DETAB-E227 and DETAB-P39 using a Qiagen minikit (Qiagen, Hilden, Germany) in accordance with the manufacturer’s instructions. Whole genome sequencing was performed using both the Illumina HiSeq (Illumina, San Diego, USA) and the Oxford Nanopore GridION (Nanopore, Oxford, UK) platforms (Tianke, Zhejiang, China). *De novo* assembly of the Illumina and Nanopore reads was performed using Unicycler v0.4.8 (Wick *et al*., 2017). MLST with the Pasteur and Oxford schemes was performed using mlst (https://github.com/tseemann/mlst) (Bartual *et al*., 2005; Diancourt *et al*., 2010).

### Plasmid characterisation

For alignment and visualisation, all plasmids were opened in the same orientation and at the same position 48 bp upstream of the GR13 *rep* gene. ARGs and *rep* genes were identified using ABRicate v0.8.13 (https://github.com/tseemann/abricate) with the ResFinder (Zankari *et al*., 2012) and pAci (Supplementary File 1) databases, respectively. Insertion sequences were identified using the ISFinder database (Siguier, 2006). To screen the entire plasmid collection, an offline version of the ISFinder nucleotide database was constructed from an available version from October 2020 (https://github.com/thanhleviet/ISfinder-sequences). The database was used with abricate, initially with a minimum nucleotide identity threshold of 80% and coverage threshold of 90% to identify putative novel IS. Representative sequences with greater than 90% coverage and between 80% and 95% nucleotide identity were validated manually and those that appeared to represent complete IS (Table S2) were added to the database. The resulting database was used with 95% identity and coverage thresholds, and sequences identified were considered isoforms of the representative IS or putative IS in accordance with ISFinder’s directions for isoform identification. Gene Construction Kit (Textco Biosoftware, Raleigh, USA) was used to annotate and examine plasmid sequences.

### Plasmid pangenome analysis

Plasmids were annotated with Prokka 1.14.6 (Seemann, 2014), using reference protein sequences to standardise annotations. Reference sequences were obtained from the NCBI Identical Protein Groups resource by querying “ `Acinetobacter[Organism] AND (uniprot[filter] OR refseq[filter])` ”. As insertion sequences were analysed separately, lines matching “transposase” or “product=IS” were removed from gff annotation files. Pangenomes and a core-gene alignment were constructed from these annotations using Panaroo 0.1.0 (Tonkin-Hill *et al*., 2020), reducing contamination-removal processes using `--mode relaxed --no_clean_edges --min_trailing_support 0 -- min_edge_support_sv 0 --trailing_recursive 0` to reflect the use of complete sequences of highly mosaic plasmids. Functional annotation based on the eggNog orthology database version 5.0.2 (Huerta-Cepas *et al*., 2019) was performed with emapper-2.1.6-43-gd6e6cdf (Cantalapiedra *et al*., 2021) using Diamond version 2.0.13 (Buchfink *et al*., 2021) for protein sequence alignments.

### Core gene analysis

Plasmid *rep* gene sequences were extracted manually, then aligned using MAFFT version 7 (Katoh *et al*., 2019) with the GNS-i iterative refinement method and additional parameters, `--reorder --anysymbol --maxiterate 2 --retree 1 –globalpair`. Low confidence residues in the alignment were masked with GUIDANCE2 (Sela *et al*., 2015). Phylogenies were constructed from the *rep* gene alignment using RaxML version 8.2.12 (Stamatakis, 2014) and the GTRGAMMA model with automated bootstrapping.

For investigation of core gene recombination, BLASTn was used to query all plasmid sequences with *parA, parB* and *tyr13* from reference plasmid p3ABAYE and identify their homologs. The resulting sequences were aligned with MAFFT as described above and a neighbour-joining phylogeny was constructed. BAPS was used to partition all core-gene phylogenies and the highest level of BAPS discrimination was used to define distinct core gene variants.

## Data availability

The complete sequences of the chromosomes and plasmids of *A. baumannii* DETAB-E227 and *A. baumannii* DETAB-B39 have been deposited in the GenBank nucleotide database under accession numbers CP073060-CP073061 and CP072526-CP072529, respectively.

## Results

### Carbapenem-resistant DETAB-E227 carries a multidrug resistance GR13-type plasmid

DETAB-E227 was resistant to imipenem, meropenem, ceftazidime, gentamicin, tobramycin and ciprofloxacin, but susceptible to colistin and tigecycline (Table S3). The complete genome of DETAB-E227 includes a 3,749,086 bp chromosome and three plasmids, pDETAB4, pDETAB5 and pDETAB6 (Table 1). DETAB-E227 is a novel sequence type according to the Pasteur (ST_IP_1554: *cpn60-3, fusA-3, gtlA-2, pyrG-79, recA-3, rplB-4, rpoB-4*) and Oxford (ST_OX_2210: *cpn60-1, gdhB-208, gltA-1, gpi-171, gyrB-231, recA-1, rpoD-153*) MLST schemes. Nine antibiotic resistance genes were found in the DETAB-E227 genome (Table 1). Two of these, *bla*_ADC-25_ and *bla*_OXA-424_, are the native AmpC and OXA-51 β-lactamase genes found in the chromosome. The *sul2* and *tet*(B) genes are in the 113,682 bp GR24-type plasmid pDETAB4 and the remaining resistance genes, *bla*_NDM-1_, *bla*_OXA-58_, *ble*_MBL_, *aacC2d* (also called *aac(3’)-IId*), *msr*(E)*-mph*(E) and a second copy of *sul2* are in the 97,035 bp GR13-type plasmid pDETAB5 (Table 1; Figure 1A).

**Table 1:** DETAB-E227 and DETAB-P39 genome characteristics

| Isolate | DNA element | Replicon type | Size (bp) | % GC | Antibiotic resistance genes |
| --- | --- | --- | --- | --- | --- |
| DETAB-E227 | chromosome | - | 3,749,086 | 39.1 | *bla*<sub>ADC-25</sub><sup>1</sup>, *bla*<sub>OXA-424</sub> |
|  | pDETAB4 | GR24 | 113,682 | 42.0 | *sul2*, *tet*(B) |
|  | pDETAB5 | GR13 | 97,035 | 41.5 | *bla*<sub>OXA-58</sub>, *bla*<sub>NDM-1</sub>, *ble*<sub>MBL</sub>, *sul2*, *aacC2d*, *msr*(E)-*mph*(E) |
| DETAB-P39 | chromosome | - | 3,877,093 | 38.9 | *bla*<sub>ADC-25</sub><sup>1</sup>, *bla*<sub>OXA-88</sub> |
|  | pDETAB13 | GR13 | 91,083 | 39.9 | - |

<sup>1</sup>The genes in DETAB-E227 and DETAB-P39 are 96.6% and 95.9% identical to the reference *bla*<sub>ADC-25</sub> sequence, respectively.

**Figure 1:**
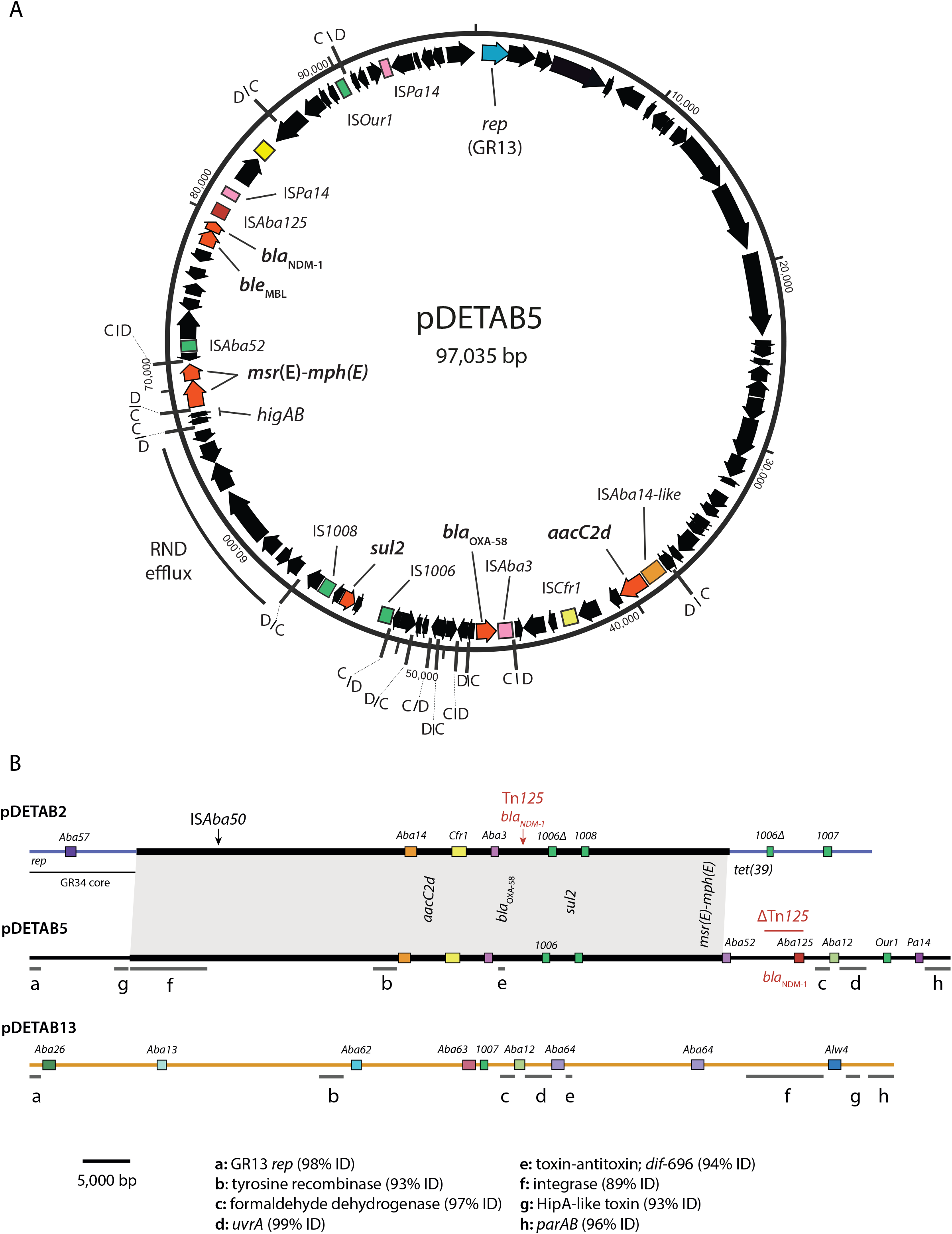
GR13 plasmid pDETAB5. **A)** Circular map of pDETAB5 drawn from GenBank accession CP072528. Plasmid sequence is shown as a black line, with arrows inside representing ORFs. Coloured boxes represent IS. Black lines marked “C/D” or “D/C” represent p*dif* sites and indicate their orientations. **B)** Linear maps of pDETAB2, pDETAB5 and pDETAB13, drawn to scale from GenBank accessions CP047975, CP072528 and CP073061. Near-identical sequences in pDETAB2 and pDETAB5 are bridged by grey shading and homologous regions of pDETAB5 and pDETAB13 are marked by lines labelled ‘a’ to ‘h’. IS are shown as coloured, labelled boxes and the locations of ARGs are indicated.

In three independent conjugation experiments, pDETAB5 transferred from DETAB-E227 to *A. baumannii* ATCC 17978 at a mean frequency of 6.96×10^−7^ transconjugants per donor (Table S4). The presence of pDETAB5 in ATCC 17978 transconjugants was confirmed using S1-PFGE, Southern blotting targeting the *bla*_NDM-1_ and *bla*_OXA-58_ genes, and PCR targeting the pDETAB5 *rep* gene (Figure S1). pDETAB5 did not transfer from DETAB-E227 to *A. nosocomialis* XH1816 or from ATCC 17978 to XH1816 in three independent experiments.

### pDETAB5 resembles the GR34 plasmid pDETAB2

The combination of ARGs in pDETAB5 resembles that in the GR34 plasmid pDETAB2, found in a ST_IP_138 *A. baumannii* isolated in the same ICU one month prior to DETAB-E227 (Liu *et al*., 2021). pDETAB5 contains 14 p*dif* sites (Figure 1A) and pDETAB2 contains 16. Alignment of pDETAB5 and pDETAB2 reveals that approximately 63 kbp of the pDETAB2 sequence is present in pDETAB5 (Figure 1B). The sequence they share includes multiple *dif* modules but not the region of pDETAB2 that has been identified as core to GR34 plasmids (Liu *et al*., 2021). The *aacC2d, bla*_OXA-58_ and *msr*(E)-*mph*(E)-containing *dif* modules in pDETAB2 and pDETAB5 are identical and their *sul2*-containing modules differ only through a 132 bp deletion in the copy of IS*1006* in pDETAB2. Other *dif* modules shared by the plasmids encode HigAB-like and AdkAB-like toxin-antitoxins, a putative serine recombinase and a putative RND efflux system.

The *bla*_NDM-1_ and *ble*_MBL_ genes in pDETAB5 and pDETAB2 are in different contexts. In pDETAB2, the *bla*_NDM-1_ and *ble*_MBL_ genes are in a complete copy of Tn*125* inserted in a 696 bp *dif* module that contains putative toxin-antitoxin genes (Liu *et al*., 2021). This module is uninterrupted in pDETAB5 (Figure 1A) and instead, the *bla*_NDM-1_ and *ble*_MBL_ genes are in a partial copy of Tn*125* that retains one copy of IS*Aba125* and 3,062 bp of the passenger segment (labelled red line in Figure 1B). This indicates that, despite sharing a collection of *dif* modules that must have a common origin, pDETAB2 and pDETAB5 acquired *bla*_NDM-1_ independently in distinct Tn*125* insertion events.

### pDETAB13 of carbapenem-sensitive DETAB-P39 is only distantly related to pDETAB5

The complete genome of DETAB-P39 includes a 3,877,093 bp chromosome and the 91,083 bp plasmid pDETAB13. DETAB-39 is ST_IP_221 and ST_OX_351. Despite growing on the initial meropenem-supplemented CHROMagar plate, DETAB-P39 was phenotypically sensitive to meropenem and to all other antibiotics tested (Table S1), and its genome does not contain any acquired antibiotic resistance genes.

The *rep* gene of pDETAB13 is 99.4% identical to that of the reference GR13 plasmid pA3ABAYE and 98.3% identical to that of pDETAB5. pDETAB13 contains eight p*dif* sites and nine complete insertion sequences (ISs), including the novel IS*Aba62*, IS*Aba63* and IS*Aba64* (Figure 1B). Excluding ISs, just 20,998 bp of pDETAB13 is homologous to pDETAB5, but the shared sequences are split across eight regions that range from 99% to 93% identical (Figure 1B). The shared segments include the *rep* gene and putative partitioning genes *parAB* in a contiguous region (a and h in Figure 1B), and a putative HipA-like toxin, a toxin-antitoxin system, formaldehyde dehydrogenase, UvrA-like excinuclease, an integrase and a tyrosine recombinase. The toxin-antitoxin genes in pDETAB13 are found in a 696 bp *dif* module (*dif*-696b) that is 93.6% identical to the one in pDETAB5 (*dif*-696a). Accumulated SNPs differentiate the *dif*-696 variants, suggesting that the presence of these modules in pDETAB5 and pDETAB13 is not the result of a recent horizontal transfer event.

Some notable ORFs in pDETAB13 are not shared by pDETAB5. A cluster of nine ORFs located in a 10,724 bp region, which we termed ADH, between IS*Aba62* and IS*Aba64* includes determinants for a putative transcriptional regulator, putative alcohol dehydrogenases and putative metabolic enzymes including a monooxygenase, amidotransferase, hydrolase, alkene reductase and oxidoreductase. A 2,111 bp *dif* module, *dif*-2111, encodes a putative NAD(P)-dependent alcohol dehydrogenase and a LysR-family transcriptional regulator. Three other *dif* modules in pDETAB13 were not assigned functions as they encode hypothetical proteins.

### GR13 plasmids have been collected from diverse sources

To characterise the GR13 plasmid family, we conducted a comparative analysis of publicly available sequences. The 1,173 bp *rep* gene of reference GR13 plasmid p3ABAYE (GenBank accession CU459140) was used to query the GenBank non-redundant nucleotide database, and 41 complete plasmid sequences containing *rep* genes greater than 74% identical to the query were found (Table S5). These GR13 plasmids are from different species, including *A. baumannii, A. pittii, A. nosocomialis, A. johnsonii, A. soli, A. seifertii* and *A. radioresistans*, and their hosts were isolated from various cities in China as well as from Japan, Cambodia, Thailand, Vietnam, India, Pakistan, Australia, Chile, the USA, the Czech Republic, France, Germany and the Netherlands between 1986 and 2020 (Table S5). Sources of isolation ranged from human clinical specimens and hospital environments to terrestrial and marine animals and environments (Table S5). The plasmids range in size from 50,047 bp to 206,659 bp and three carry additional replication genes (Table S5), suggesting that they have formed cointegrates with plasmids from different *rep* groups.

### The small GR13 core genome has been subject to recombination

To characterise the gene content of GR13 plasmids, a pangenome was constructed. This consisted of 1190 genes: two considered core (present in 43 plasmids), two soft-core (42 plasmids), 123 shell (7 to 40 plasmids) and 1060 cloud (1 to 6 plasmids).

The four core genes were *rep*, the putative partitioning genes *parA* and *parB*, and a putative tyrosine recombinase gene that we will refer to as *tyr13*. Though *parAB* were found in only 42 of the 43 plasmids by the pangenome approach, using tBLASTn to query the remaining plasmid sequence (CP038259) with the amino acid sequences of ParA and ParB of pDETAB13 revealed equivalent genes with nucleotide identities of 78.8% and 79.8%, respectively. The *parAB* genes were found adjacent to one another in all 43 plasmids and were usually adjacent to *rep*, but *tyr13* was never found adjacent to *rep* or *parAB*.

The conservation of core genes was investigated by using BAPS to place gene sequences into allelic groups that differed by few SNPs and exhibited common SNP patterns that likely arose cumulatively from a recent ancestor. The distribution of *parAB* and *tyr13* allelic groups were visualised relative to a *rep* gene phylogeny (Figure 2A) and instances of recombination were identified where phylogenetic clusters did not contain *parA, parB* and *tyr13* from the same allelic groups. Substitution of *tyr13* genes appears to have occurred on multiple occasions while a single example of *parB* allele substitution was seen in CP022299.

**Figure 2:**
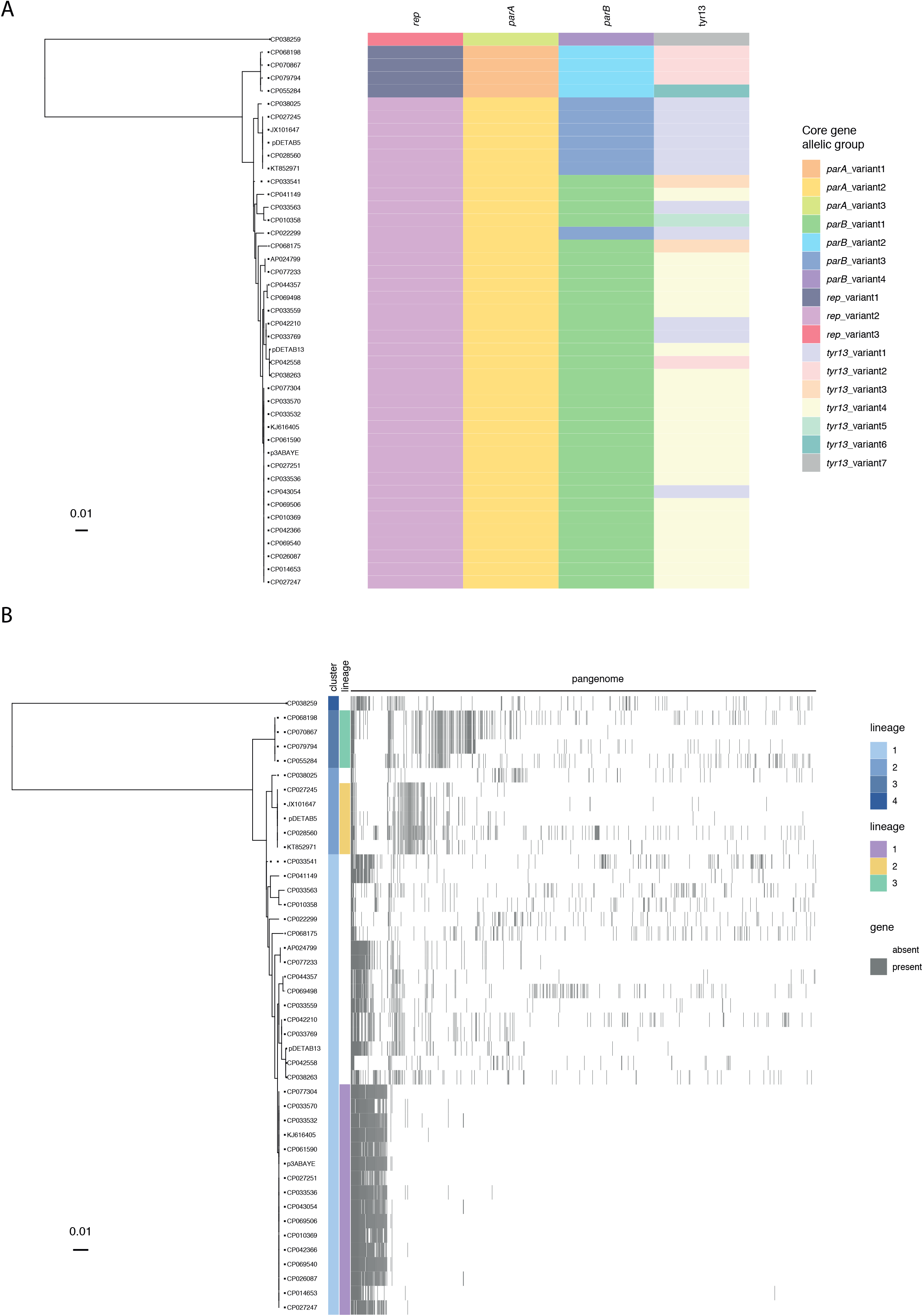
The GR13 plasmid family pangenome. **A)** Plasmid core gene allelic group identities displayed relative to a *rep* gene phylogeny. **B)** GR13 pangenome displayed relative to the *rep* gene phylogeny. Cluster and lineage memberships are indicated to the left of the pangenome visualisation.

### GR13 plasmid lineages have disseminated widely

The *rep* gene phylogeny was used to sub-type GR13 plasmids. The collection was partitioned into four broad-ancestry clusters of plasmids that, apart from CP022299, shared core genes from the same allelic groups, reflecting their common ancestry. Further *rep* gene variation, as evident in the phylogenetic tree (Figure 2B), indicated that clusters could be partitioned into epidemiologically relevant plasmid lineages. We have defined three GR13 lineages that are represented by four or more plasmids in this collection that are not separated by any SNPs in the *rep* gene phylogeny (Figure S2), equivalent to a total *rep* identity of >99.8% for lineage 1 and 100% for lineages 2 and 3. Plasmids in the same lineage share significant accessory gene content (Figure 2B), consistent with them having descended from an ancestral plasmid that contained the same *rep* gene and a conserved set of accessory genes. The presence of accessory genes that differentiate some plasmids from other members of the same lineage highlight their capacity to diversify through gene acquisition.

Host species and sources of isolation varied within lineages, indicating that they have disseminated internationally and between *Acinetobacter* species. The best-represented lineage in this collection, lineage 1, contains the reference GR13 plasmid p3ABAYE and 15 others. p3ABAYE is from a clinical *A. baumannii* isolated in France in 2001, while other members of lineage 1 are from *A. pittii, A. nosocomialis* and *A. seifertii* strains from human clinical samples in various Chinese provinces, Australia, Colombia and Germany (Table S5). A single lineage 1 plasmid is derived from marine sediment. Lineage 1 has a well-conserved accessory genome, consisting of 39 core genes (present in all 16 plasmids), 58 shell genes (in two to 15 plasmids) and just two cloud genes (in one plasmid each). Lineage 2 plasmids include pDETAB5 and four other ARG-bearing plasmids. Representatives of lineage 2 have been found in *A. baumannii, A. soli* and an isolate of indeterminate *Acinetobacter* species derived from clinical samples, an ICU environment or sewage, but only in mainland China, Taiwan or Vietnam. In contrast, the four representatives of lineage 3 are from *A. johnsonii* or an indeterminate *Acinetobacter* that were isolated across wide geographic and temporal spans: soil from the USA in 1986, a spacecraft-associated clean room in the Netherlands in 2008, an intensive care unit sink in Pakistan in 2016 and bigeye tuna in China in 2018. Taken together, the distributions of lineage 1, 2 and 3 plasmids emphasise the capacity of GR13 plasmids for widespread dissemination and persistence.

### Most accessory genes in GR13 plasmids are unique

There were 1090 gene families in the GR13 pangenome, with 4453 genes identified in total. Of the 1090 gene families, 745 could be assigned putative functions with our Prokka annotation and Panaroo approach (62.6%), while 526 (44.2%) and 442 (37.1%) were assigned functions with the COG and KEGG schemes, respectively. COG placed gene families in broad functional categories, most commonly replication and repair (135/526, 25.7%), transcription (83, 15.8%) and inorganic transport and metabolism (78, 14.8%) (Supplementary File 2). KEGG categories offered more specific functional annotation and facilitated identification of the most common gene functions in the collection. Amongst the 50 most prevalent gene families that were assigned functions (Supplementary File 2), families with putative metabolic functions were most common. The second most common gene families encode components of toxin-antitoxin systems, with HipA-like and Fic-like toxin genes the most abundant overall. Other functions of note included DNA integration and recombination (42 gene families, 194 genes, 43 plasmids), antimicrobial resistance (3 gene families, 6 genes, 5 plasmids), heavy metal resistance (22 gene families, 51 genes, 7 plasmids), alcohol tolerance (11 gene families, 172 genes, 32 plasmids), and phage defence (5 gene families, 89 genes, 17 plasmids). Some functional groups contained multiple gene families, suggesting that genes with the same functions have been acquired on multiple occasions by different GR13 lineages.

Of the 1186 accessory gene families, 746 (62.9%) were found in just a single plasmid each. These so-called “singleton genes” were found in 25 of the 43 plasmids, where they accounted for between 0.7% and 61.4% of total gene content. The pangenome network showed that many singleton genes were found adjacent to one another in long, contiguous sequences that were unique to the plasmids that carried them (Supplementary File 3). In CP028560 and CP069498, an abundance of singleton genes coincided with the presence of additional *rep* genes of types GR34 and GR24, respectively, indicating that plasmid cointegrate formation was associated with the introduction of significant numbers of novel genes.

### ARG-bearing dif modules are subject to rearrangement by insertion sequences

Lineage 2 plasmids and the only other ARG-containing plasmid, CP033563 from an *A. nosocomialis* isolated in Taiwan in 2010, carry ARGs in *dif* modules. Only two plasmids from lineage 2 appear to have acquired additional ARGs: KT852971 has acquired a *sul1*-containing class 1 integron with cassette array *aadB*-*arr-2*-*cmlA1*-*aadA1* and JX101647 has acquired *sul2* and *aphA1* in an insertion within an existing ARG-containing *dif* module, described below.

To assess whether and how individual *dif* modules have evolved since they were acquired by the ancestor of lineage 2 plasmids, their ARG-containing *dif* modules were compared. The *msr*(E)-*mph*(E) module was unchanged between the four plasmids that carried it, but variation was seen across *bla*_OXA-58_ (Figure 3A), *sul2* (Figure 3B) and *aacC2d*-containing (Figure 3C) modules. The novel IS elements IS*Aso1* and IS*Aso2*, members of the uncharacterised IS*NCY* family, were acquired by both the *bla*_OXA-58_ and *sul2-*containing modules in JX101647 (Figure 3). Both IS inserted in the same orientation adjacent to the XerC binding ends of p*dif* sites that flank their respective *dif* modules (Figure 3A and B), with the 5 bp immediately adjacent to p*dif* involved in the target site duplication generated by insertion. The IS*6*/IS*26*-family element IS*1008* fused the *bla*_OXA-58_-containing module of CP027245 to the remnant of a previously described RND efflux module (Liu *et al*., 2021) following a deletion of indeterminate length (Figure 3A). IS*1008* also deleted part of the *sul2-*containing module of CP028560 and brought the remainder adjacent to another sequence, possibly a remnant of a *floR*-containing module as IS*1008* has truncated the *floR* gene (Figure 3B). In JX101467 a partial deletion of the IS*Aba14*-like element is associated with the acquisition of a 12,213 bp segment bounded at one end by an IS*Our1*-like element and at the other by an IS*Alw27*-like element (Figure 3C). The acquired segment includes *sul2* and a truncated copy of Tn*5393* that is interrupted by the *aphA1*-containing Tn*4352*. These examples highlight the capacity of IS to influence the accessory content of *dif* modules through insertion and by mediating deletion events.

**Figure 3:**
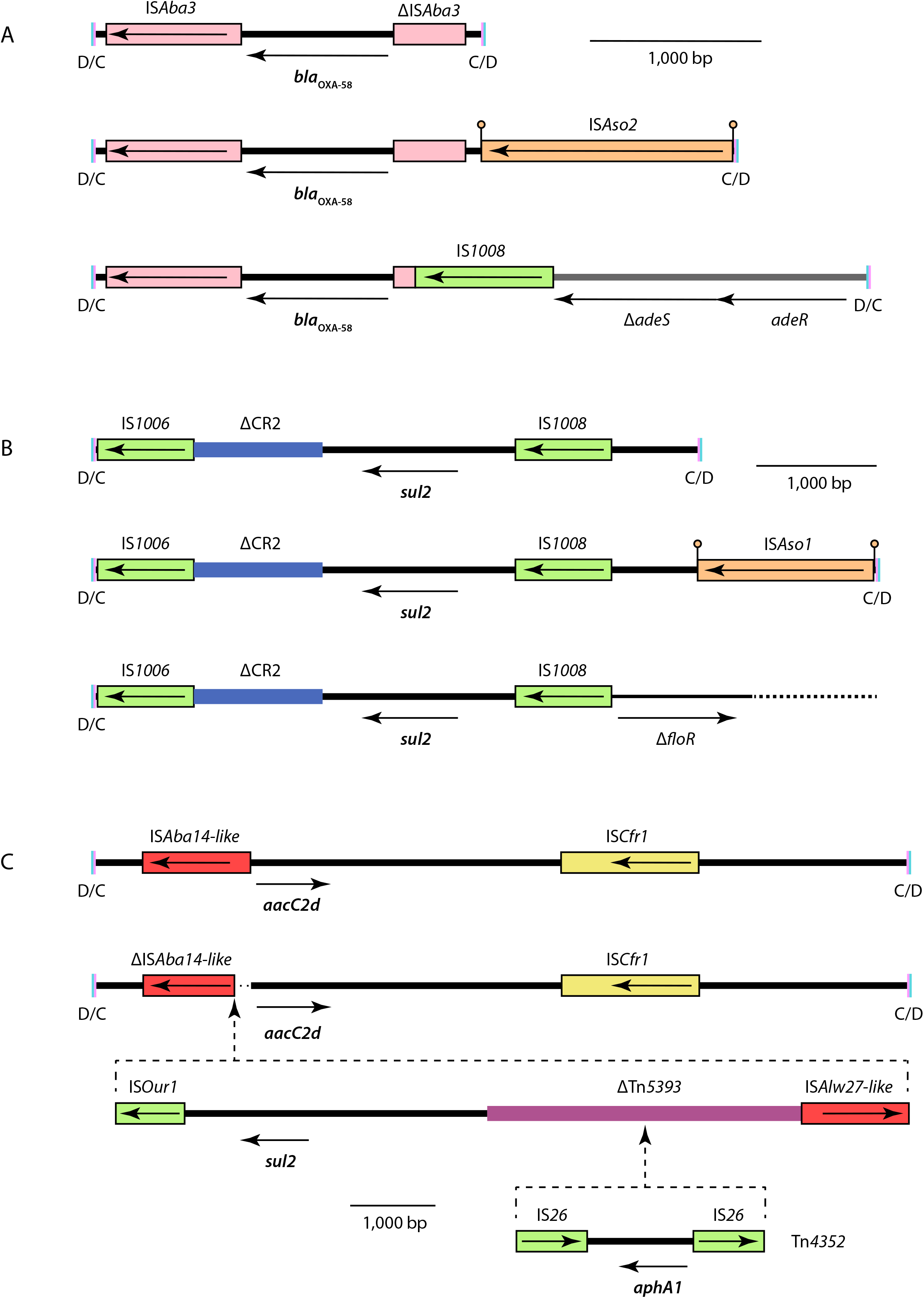
ARG-containing *dif* module variants. Scaled diagrams of *dif* modules containing **A)** *bla*_OXA-58_, **B)** *sul2*, and **C)** *aacC2d*. The extents and orientations of ORFs are indicated by labelled horizontal arrows and IS are shown as labelled boxes. IS that are the same colour belong to the same family. Drawn to scale from GenBank accessions CP072528, JX101647, CP028560, KT852971 and CP027245.

### Diverse accessory genes are found in dif modules

To further characterise their potential for mobilising accessory genes other than the well-known ARGs, we examined the content and distribution of 17 *dif* modules identified in pDETAB5, pDETAB13, p3ABAYE, AP024799, CP022299 and CP068175 (Table S6). The sequences of these modules were used to query the GR13 collection with BLASTn and their distributions are shown in Figure 4A. Twelve *dif* modules were only carried by the plasmid or plasmid lineage that they were identified in, but five were found in multiple lineages, suggesting that they have been acquired independently.

**Figure 4:**
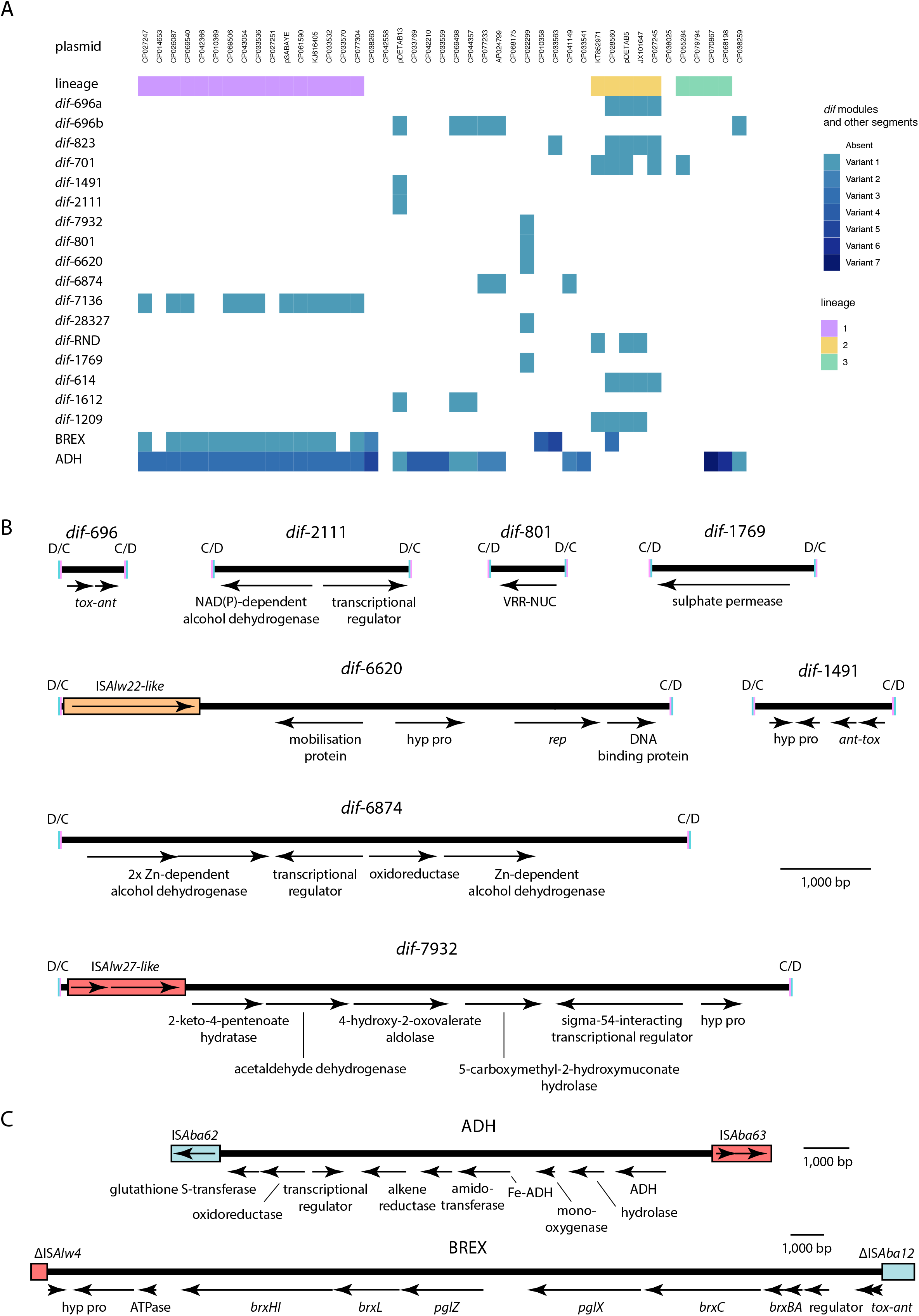
Novel *dif* modules carrying diverse accessory genes. **A)** Presence and absence of *dif* modules identified in GR13 plasmids. Plasmids are ordered as in the *rep* gene phylogeny shown in Figure 2, and membership of lineages 1, 2 and 3 is indicated. The presence of variant BREX and ADH segments are indicated by different shades of colour B) Scaled diagrams of selected *dif* modules. **(C)** Scaled diagrams of accessory gene segments. Sequences in parts B and C were drawn to scale from GenBank accessions CP072528, CP073061, CU459140, AP024799, CP022299 and CP068175.

Three dif modules identified here (*dif*-2111, *dif*-6874 and *dif*-7136) encode putative alcohol dehydrogenases which may be involved with alcohol tolerance and quorum sensing (Lin *et al*., 2021). The largest module we identified, *dif*-28327, encodes putative copper resistance proteins, and *dif*-7932 encodes a set of putative metabolic proteins that appear to be involved with aromatic compound degradation (Figure 4B). The *dif*-1769 module carries a putative sulphate permease determinant and is 88% identical to part of a *sulP* module that has been described previously (Mindlin *et al*., 2018). A module found only in the cointegrate plasmid CP022299 contains a *rep* gene 82.3% identical to the reference GR26 *rep* (GenBank accession CP015365), as well as a putative mobilisation gene (Figure 4B). This appears to be a small plasmid that has been integrated through recombination at p*dif* sites. The remaining modules could not be assigned putative functions, though one of these, *dif*-801, encodes a protein with a VRR-NUC domain (Pfam: PF08774), equivalents to which have been described in restriction endonucleases (Kinch *et al*, 2005).

Two sets of ORFs from the GR13 shell genome that appeared to be discrete units in the genome network were examined to determine whether they were found in well-conserved *dif* modules. The first set of ORFs resemble determinants for bacteriophage exclusion (BREX) systems (Goldfarb *et al*., 2015), and likely have anti-phage functions. The BREX determinants are not part of an identifiable *dif* module. Instead, they are found in a 26,140 bp segment flanked by partial copies of IS*Alw4* and IS*Aba12* (Figure 4C). The same BREX segment is present in 14 of 16 plasmids from lineage 1, while variant sequences are present in four plasmids from elsewhere in the phylogeny (Figure 4A). The second set of well-conserved ORFs correspond to the ADH segment of pDETAB13 that contains two putative alcohol dehydrogenase determinants, as well as several ORFs with expected metabolic functions and one for a putative transcriptional regulator (Figure 4C). Variants of the ADH segment are found in 30 of the 43 GR13 plasmids studied here (Figure 4A).

### Diverse insertion sequences shape GR13 plasmid accessory content

We used a version of the ISFinder database to screen the GR13 plasmid collection and assess the abundance, diversity and richness of IS. Individual plasmids contained between one and 34 IS, with up to 18 different IS and up to six copies of the same IS found in a single plasmid (Figure S3). Seventy-five different IS were found, representing 15 different IS families. These included 26 putatively novel IS that differed from named sequences by greater than 5% nucleotide identity. Five of these, including the three identified in pDETAB13, were characterised, submitted to ISFinder and named as part of this study (Figure S3).

Members of the IS*3* (22 different IS) and IS*5* (15 different IS) families dominated the IS population, with representatives of one or both found in all but two GR13 plasmids. The next best-represented family was IS*6*/IS*26* (8 different IS), members of which are known to be strongly associated with antibiotic resistance (Harmer and Hall, 2019). The highest numbers of IS*6*/IS*26* family elements were found in the ARG-containing lineage 2 plasmids, where many were associated with ARG-containing *dif* modules (green in Figure 3), but these elements were also seen in 13 plasmids that do not contain ARGs. IS*NCY*-family IS (7 different IS; 26 copies) including IS*Alw2*2, IS*Aso1*, IS*Aso2* (Figures 3 and 4B), and four putative IS identified here are distributed throughout the GR13 family.

## Discussion

Our discovery of two GR13-type plasmids in unrelated *A. baumannii* strains isolated a month apart in the same ICU, one cryptic and the other conferring multi-drug resistance, prompted an investigation of the wider GR13 plasmid family, which had not been studied previously. GR13 plasmids are found in multiple *Acinetobacter* species from a diverse set of environments. The four-gene core of GR13 plasmids is associated with a diverse accessory genome influenced by the exchange of *dif* modules, the acquisition of translocatable elements, and IS-mediated deletions. This characterisation of a family of *Acinetobacter* plasmids that can carry clinically-significant ARGs adds to a growing body of literature on the accessory genepool of this important human pathogen and the underestimated role plasmids in generating diversity across this genus.

### Diversity in GR13 plasmids: consequences for genomic surveillance and epidemiology

Here, we identified three GR13 lineages on the basis of *rep* gene identity that we found share lineage-specific sets of accessory genes. Although *rep* or core-gene typing cannot account for all accessory gene diversity within GR13 lineages, we found that plasmids in the same lineage share significant gene content. Lineage-specific markers like *rep* and *parAB* might be used in targeted surveillance programs to detect clinically-relevant plasmids such as pDETAB5 and other members of lineage 2. Representatives of lineage 2 have so far only been seen in isolates from China or neighbouring Vietnam (Table S2), where the first example appeared in 2005, but it will be interesting to trace this lineage and monitor the dynamics of its dispersal in epidemiological studies. We have provided the sequences of the *rep* and *parAB* genes that can be used to identify plasmids from lineages 1, 2 and 3 in Supplementary File 4. These can be used for higher-resolution genomic surveillance to track the dissemination of GR13 lineages across *Acinetobacter*.

### How do GR13 plasmids spread horizontally?

Plasmids from the same GR13 lineages have been found in different host species that have been isolated from various sources and geographic locations. This is clear evidence for their widespread dissemination and ability to replicate in various *Acinetobacter* species. However, the mechanisms responsible for the horizontal transmission of GR13 plasmids remain unclear. In this study, pDETAB5 transferred from DETAB-E227 to *A. baumannii* ATCC 17978 at a relatively low frequency, but failed to transfer from DETAB-E227 or ATCC 17978 to *A. nosocomialis* strain XH1816.

No candidate set of ORFs for a type IV secretion system that might be associated with conjugation were found in pDETAB5 or any of the GR13 plasmids examined here, so it appears they rely on alternative mechanisms for horizontal transfer. In contrast, other large *Acinetobacter* plasmids have been shown to carry conjugation determinants in conserved backbones (Hamidian *et al*., 2016; Nigro *et al*., 2014) while small plasmids that carry an origin-of-transfer (*oriT*) and cognate mobilisation genes (Hamidian and Hall, 2018) or *oriT* alone (Blackwell and Hall, 2019) can be mobilised by co-resident conjugative plasmids. It is possible that the integration of small mobilisable plasmids through recombination at p*dif* sites contributes to the mobility of GR13 plasmids. An example of this is seen in CP022299 where all or part of a putatively mobilisable plasmid is present in the *dif*-6620 module (Figure 4). The acquisition of *oriT* sequences through small plasmid integration has been described for large plasmids in *Staphylococcus* and *Proteus* (Hua *et al*., 2020; O’Brien *et al*., 2015), though in those cases integration did not involve p*dif* sites. Horizontal transfer in outer membrane vesicles has also been reported in *Acinetobacter* (Chatterjee *et al*., 2017; Rumbo *et al*., 2011) and this, or other passive DNA transfer mechanisms, might play a role in plasmid dispersal.

### Site-specific recombination and tyrosine recombinase genes in GR13 plasmids

The importance of site-specific recombination to the evolution of some types of plasmids in *Acinetobacter* has become increasingly evident. XerC and XerD-mediated recombination at p*dif* sites is implicated in the movement of *dif* modules between plasmids of different types (Blackwell and Hall, 2017; Hamidian *et al*., 2021; Hamidian and Hall, 2018), and has been shown experimentally to generate cointegrate plasmids (Cameranesi *et al*., 2018). Recombination at p*dif* sites can also resolve cointegrates, potentially generating hybrids of the initial cointegrate-forming molecules (Cameranesi *et al*., 2018). This process likely explains the striking similarity of the GR13 plasmid pDETAB5 and GR34 plasmid pDETAB2 (Figure 1B). Supporting this hypothesis, another plasmid examined here, CP028560, is a cointegrate with GR13 and GR34 replicons identical to those in pDETAB5 and pDETAB2, and appears to represent an evolutionary intermediate.

Given p*dif* sites appear to play a major role in the evolution of some plasmids, it will be important to define the types of plasmids that carry them and can participate in XerC/D-mediated cointegration events or the exchange of *dif* modules. A recently characterised family of *Acinetobacter* plasmids has pangenome characteristics similar to the GR13 family, and representatives carry mosaic regions that were called ‘hotspots’ (Ghaly *et al*., 2020). The movement of *dif* modules might explain the dynamics of these hotspot regions. It will be useful to identify and study specific p*dif*-containing plasmid lineages over sustained periods of time to track small-scale evolutionary changes and further our understanding of the evolutionary consequences of p*dif* carriage.

### The *dif* module gene repertoire continues to grow

The first-described *dif* modules contained ARGs, but further studies have revealed that these mobile elements can carry a diverse array of passenger genes. Our characterisation of selected *dif* modules in GR13 plasmids expands the known repertoire of genes associated with these elements, further highlighting their important role in the diversification of the *Acinetobacter* accessory genome.

Many modules with diverse functions, including those expected to contribute to clinically-relevant traits such as antibiotic resistance or alcohol tolerance, are accompanied by one or more *dif* modules carrying toxin-antitoxin genes (Blackwell and Hall, 2017; Hamidian and Hall, 2018; Liu *et al*., 2021). These are expected to contribute to the stability of their host plasmids, and therefore to co-resident *dif* modules. ORFs with toxin-antitoxin functions made up 16% of functionally-annotated gene families in GR13 plasmids, suggesting that they play an important role in plasmid persistence. Diversity seen in toxin-antitoxin modules here and elsewhere support the hypothesis that these and other *dif* modules are ancient elements that have co-evolved with the plasmids of *Acinetobacter* (Hamidian and Hall, 2018).

### Insertion sequences target and reshape dif module-containing plasmids

By definition IS do not encode proteins other than those required for their transposition, but their actions can profoundly influence the evolution of their host molecules (Vandecraen *et al*., 2017). In this study we found cases where IS that are expected to generate target site duplications on insertion are not flanked by them, suggesting that they have mediated deletion events. These deletion events have clearly been responsible for sequence loss from *dif* modules, or the fusion of *dif* modules to other sequences (Figure 3). It appears IS-mediated deletion events can produce novel, hybrid *dif* modules, though whether these are mobile is likely to depend on the precise sequences of their new flanking p*dif* sites (Hamidian *et al*., 2021). IS-mediated deletions might also remove p*dif* sites associated with *dif* modules, creating larger segments that might resemble the IS-flanked BREX and ADH segments (Figure 4C).

Two elements characterised here, IS*Aso1* and IS*Aso2*, are distantly related to one another (encoding 71.0% identical transposases), but inserted in the same orientation immediately adjacent to the XerC binding regions of p*dif* sites (Figure 3). Together with previous descriptions of related IS (Blackwell and Hall, 2017; Hamidian and Hall, 2018), our findings support the notion that this group of IS*Ajo2*-like elements are “*dif* site hunters”. The presence of *dif* site hunters can be considered strongly indicative of the presence of p*dif* sites in *Acinetobacter* plasmids, and might aid in the identification of plasmid types that participate in the exchange of *dif* modules.

## Conclusions

GR13 plasmids have the capacity to accumulate diverse accessory sequences that may provide fitness advantages in the wide array of environments inhabited by *Acinetobacter* species. Some accessory modules pose risks to human health and might contribute to the persistence of *Acinetobacter* populations in hospital environments. GR13 plasmid lineages have disseminated internationally and amongst different *Acinetobacter* species. Genomic surveillance should be coupled with experimental characterisation of these plasmids to better understand their contribution to the diversification and success of *Acinetobacter*, particularly in nosocomial settings.

## Acknowledgements

We are grateful to the doctors and nurses in the ICU for sample collection. We thank Prof. Zhiyong Zong and his team for their careful teaching of sampling methods.

## Supplementary figure legends, tables and files

**Figure S1:** Transfer of pDETAB5. A) *Apa*I-treated genomic DNA from DETAB-E227, ATCC 17978 and putative ATCC 17978 transconjugants after pulsed-field gel electrophoresis. (PFGE) B) Agarose gel showing the products of PCRs targeting the *rep* genes of pDETAB4 (GR24) and pDETAB5 (GR13). The source of template DNA for each reaction is labelled above, with (-) indicative of a no-DNA control. The sizes in base pairs of labelled DNA size marker bands are indicated to the left. C) S1-treated DNA after PFGE and hybridisation with *bla*_NDM-1_ and *bla*_OXA-58_-specific probes. The sizes of bands in the DNA size marker (in kilobase pairs) are indicated to the left.

**Figure S2**: GR13 *rep* gene SNP matrix. The numbers of SNPs between *rep* genes and plasmid lineage memberships are indicated by shading as outlined in the legends to the right of the grid.

**Figure S3:** Insertion sequences in GR13 plasmids. Presence/absence matrix for insertion sequences in plasmids ordered according to the *rep* gene phylogeny. Shading indicates IS presence, with the degree of shading reflective of IS copy number as shown in the legend to the right. The colour of shading reflects IS family membership. IS with names in black text have been characterised previously, while those with names in red were characterised as part of this study. The names of putative IS identified here are grey.

**Table S1: Primers used to detect DETAB-E227 plasmid replicons via PCR**

**Table S2: Putative novel IS found in the GR13 plasmid collection**

**Table S3: Antibiotic minimum inhibitory concentrations**

**Table S4: Transfer frequency of pDETAB5 from DETAB-E227 to ATCC 17978**

**Table S5: GR13 plasmids in GenBank**

**Table S6: Characteristics of the examined dif modules**.

**Supplementary File 1. FASTA file representing the pAci database**. The pAci database includes *rep* genes of *Acinetobacter* plasmids. Plasmid types without identifiable *rep* genes are represented by other backbone genes (e.g. partitioning or transfer genes).

**Supplementary File 2. Functional annotation of the GR13 plasmids pangenome**.

**Supplementary File 3. Network visualisation of the GR13 plasmids pangenome**. This .gml file can be opened using Cytoscape (https://cytoscape.org/).

**Supplementary File 4**. *FASTA file with lineage-specific rep and parAB markers for GR13 plasmids*

