## Supplementary figures and images for "GR13-type plasmids in *Acinetobacter* potentiate the accumulation and horizontal transfer of diverse accessory genes"

### Figure S1

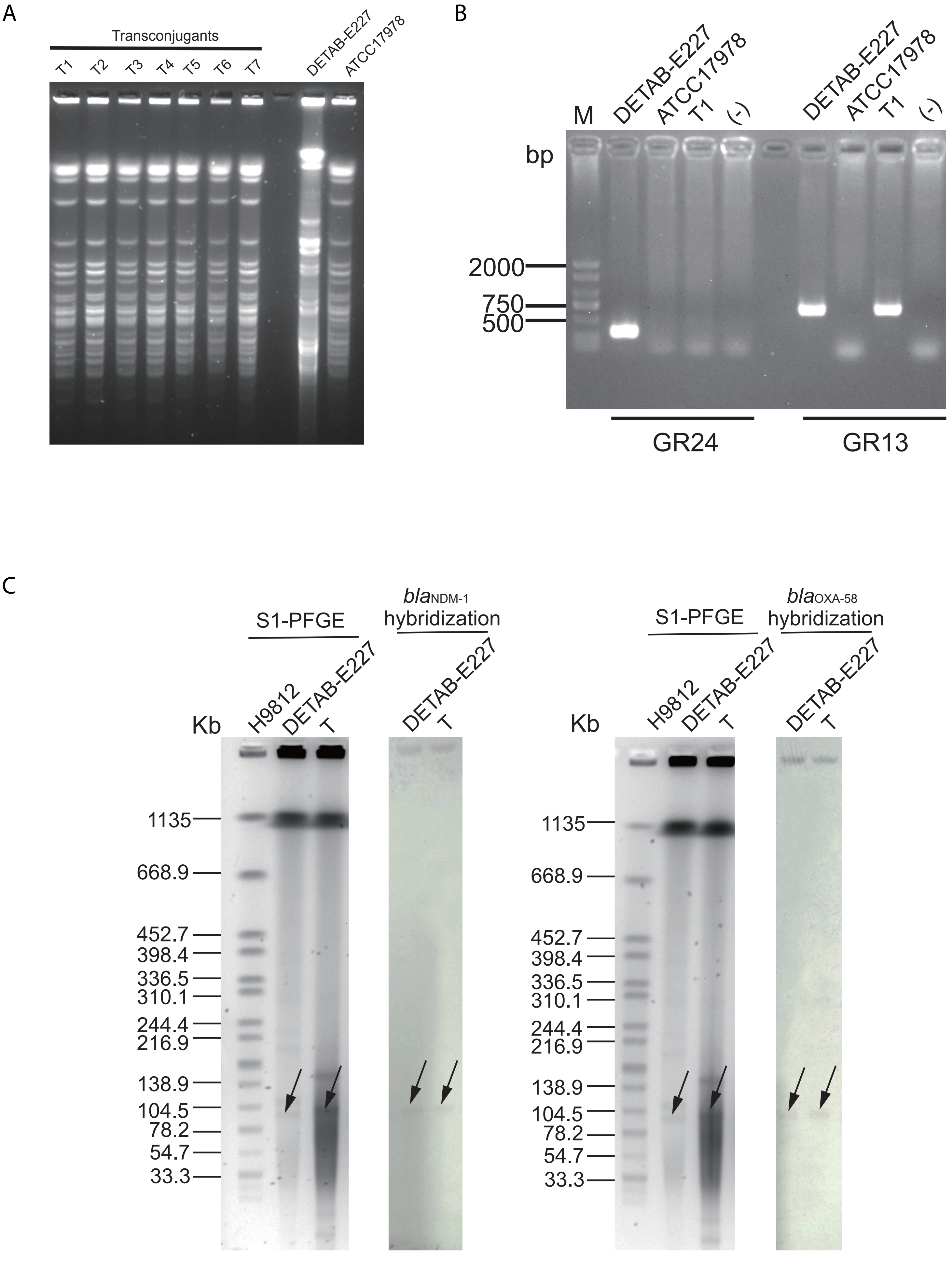

### Figure S2

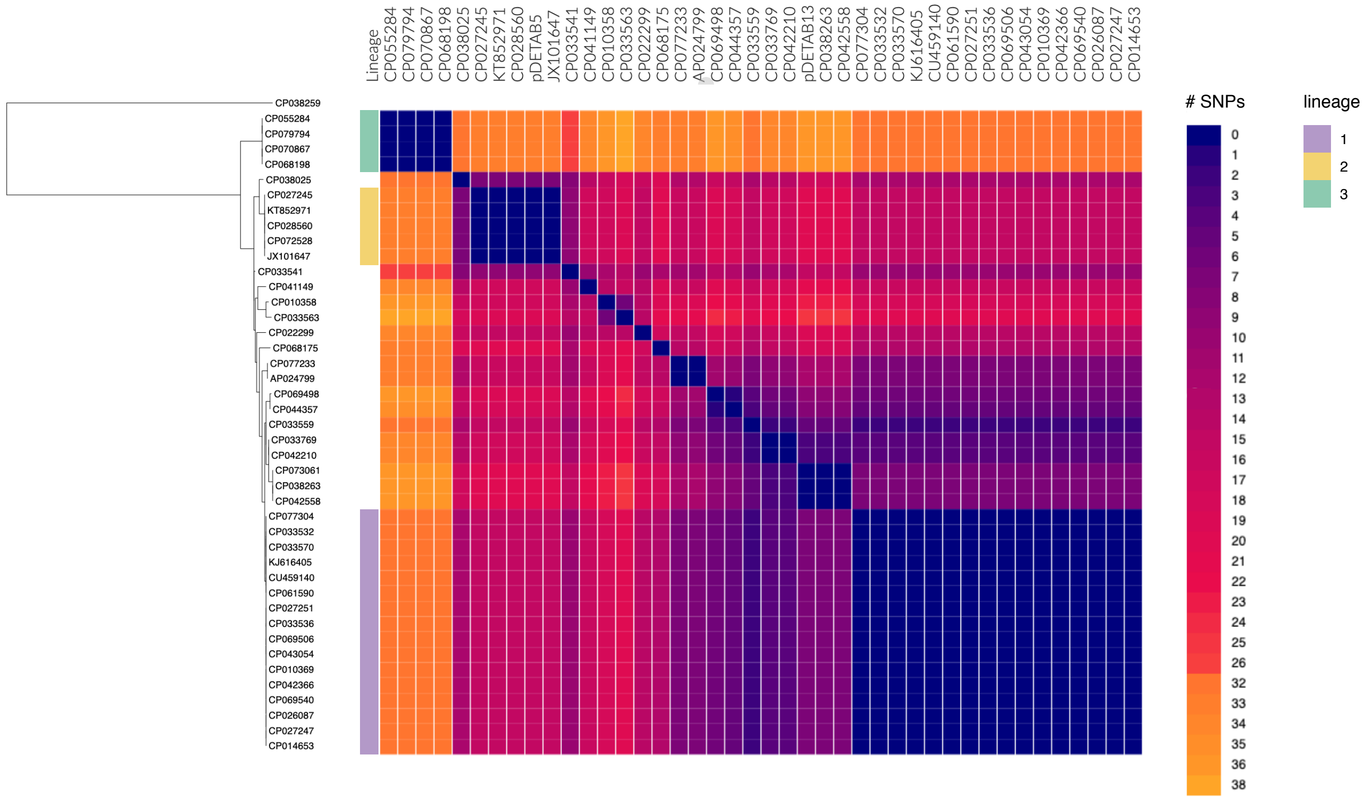

### Figure S3

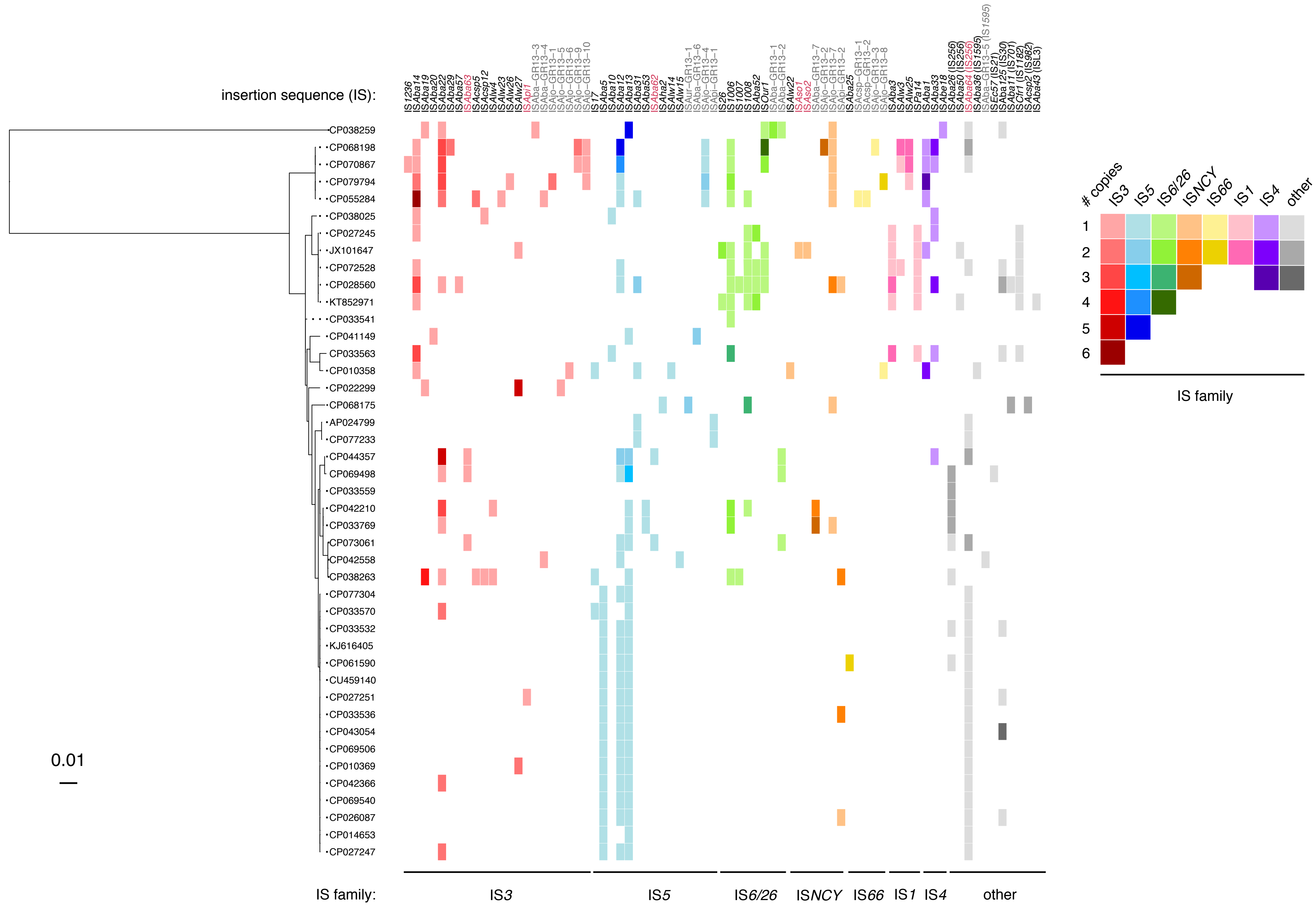
