## Supplementary material for "GR13-type plasmids in *Acinetobacter* potentiate the accumulation and horizontal transfer of diverse accessory genes": Table S1

**Table S1:** Primers used to detect DETAB-E227 plasmid replicons via PCR

| Replicon type | Plasmid | Primer name | Primer sequence (5’-3’) | Expected product  size (bp) |
| --- | --- | --- | --- | --- |
| GR24 | pDETAB4 | Hgz_103-F  Hgz_103-R | TGGCAAGATTGAGGTGGTTC  AAGTTGGTCATATCCGTACTTTCG | 327 |
| GR13 | pDETAB5 | GR13-F  GR13-R | TAGTAACCGTCTGATTAGAC  GACCTTTCTTGATGGTATCG | 680 |

All primers were used in PCRs with annealing temperatures of 55°C.
