## Supplementary material for "GR13-type plasmids in *Acinetobacter* potentiate the accumulation and horizontal transfer of diverse accessory genes": Table S2

**Table S2:** Putative novel IS found in the GR13 plasmid collection

| Sequence | Closest match (family) | % ID | Plasmid |
| --- | --- | --- | --- |
| ISAba-GR13-1 | IS*1006* (IS*6*/*26*) | 94.9 | CP038259 |
| ISAba-GR13-2 | IS1007 (IS*6*/*26*) | 94.9 | CP038259 |
| ISAba-GR13-3 | IS*1236* (IS*3*) | 93.0 | CP038259 |
| IS*Aba62* | IS*Aba12* (IS*5*) | 92.6 | pDETAB13 |
| ISAur-GR13-1 | IS*Aba12* (IS*5*) | 94.8 | CP068175 |
| ISAcsp-GR13-1 | IS*Aba17* (IS*66*) | 88.9 | CP055284 |
| ISAjo-GR13-1 | IS*Aba19* (IS*3*) | 94.4 | CP079794 |
| IS*Aba64* | IS*Aba26* (IS*256*) | 86.2 | pDETAB13 |
| ISAba-GR13-4 | IS*Aba29* (IS*3*) | 86.7 | CP042558 |
| ISAjo-GR13-2 | IS*Aba32* (IS*NCY*) | 92.8 | CP068198 |
| ISAjo-GR13-3 | IS*Aba46* (IS*66*) | 93.8 | CP068198 |
| ISAcsp-GR13-2 | IS*Aba49* (IS*66*) | 93.7 | CP055284 |
| ISApi-GR13-1 | IS*Aba5* (IS*5*) | 92.2 | AP024799 |
| ISAjo-GR13-4 | IS*Aba62* (IS*5*) | 89.0 | CP079794 |
| ISAjo-GR13-5 | IS*Aba63* (IS*3*) | 94.5 | CP022299 |
| ISAjo-GR13-6 | IS*Aca1* (IS*3*) | 90.8 | CP010358 |
| ISAba-GR13-5 | IS*Acra1* (IS*1595*) | 88.0 | CP042558 |
| IS*Api1* | IS*Acsp3* (IS*3*) | 94.2 | CP027251 |
| ISAba-GR13-6 | IS*Aha1* (IS*5*) | 92.3 | CP041149 |
| ISAjo-GR13-7 | IS*Ajo2* (IS*NCY*) | 91.6 | CP068198 |
| ISAjo-GR13-8 | IS*Alw16* (IS*66*) | 94.2 | CP079794 |
| ISAba-GR13-7 | IS*Alw22* (IS*NCY*) | 91.0 | CP033769 |
| IS*Aba63* | IS*Alw4* (IS*3*) | 94.5 | pDETAB13 |
| ISAjo-GR13-9 | IS*Alw4* (IS*3*) | 94.7 | CP068198 |
| ISAjo-GR13-10 | IS*Alw5* (IS*3*) | 92.9 | CP068198 |
| ISApi-GR13-2 | IS*Ajo2* (IS*NCY*) | 89.7 | CP033536 |
| IS*Aso2* | IS*Ajo2* (IS*NCY*) | 93.5 | JX101647 |
| IS*Aso1* | IS*Alw22* (IS*NCY*) | 89.3 | JX101647 |
