## Supplementary material for "GR13-type plasmids in *Acinetobacter* potentiate the accumulation and horizontal transfer of diverse accessory genes": Table S3

**Table S3:** Antibiotic minimum inhibitory concentrations

| Isolates | Antibiotic^1^ minimum inhibitory concentration (mg/L)^2^ | | | | | | | | |
| --- | --- | --- | --- | --- | --- | --- | --- | --- | --- |
|  | IMP | MEM | CAZ | GEN | TOB | LEV | CIP | COL | TGC |
| DETAB-P39 | 0.125  (S) | 0.5  (S) | 4  (S) | 0.5  (S) | 1  (S) | 0.06  (S) | 0.125  (S) | 1  (S) | 0.25  (S) |
| DETAB-E227 | 256  (R) | 128  (R) | >256  (R) | 256  (R) | 16  (R) | 4  (I) | 4  (R) | 1  (S) | 0.5  (S) |
| ATCC17978 | 0.5  (S) | 0.25  (S) | 4  (S) | 0.5  (S) | 0.5  (S) | 0.125  (S) | 0.125  (S) | 0.5  (S) | 0.5  (S) |
| ATCC17978  (pDETAB5)^3^ | 256  (R) | 256  (R) | >256  (R) | >256  (R) | 8  (I) | 0.125  (S) | 0.125  (S) | 0.5  (S) | 0.25  (S) |

^1^ IMP = imipenem, MEM = meropenem, CAZ = ceftazidime, GEN = gentamicin, TOB = tobramycin, LEV = levofloxacin, CIP = ciprofloxacin, COL = colistin, TGC = tigecycline

^2^ minimum inhibitory concentrations classed as resistant (R), intermediate (I) or sensitive (S)

^3^ transconjugant derived from mating DETAB-E227 with ATCC 17978
