## Supplementary material for "GR13-type plasmids in *Acinetobacter* potentiate the accumulation and horizontal transfer of diverse accessory genes": Table S4

**Table S4:** Transfer frequency of pDETAB5 from DETAB-E227 to ATCC 17978

|  | Experiment | | | |
| --- | --- | --- | --- | --- |
|  | 1 | 2 | 3 | Mean |
| #Donors (D) | 1.90×10^9^ | 1.80×10^9^ | 4.2×10^9^ |  |
| #Transconjugants (TC) | 1.75×10^3^ | 1.65×10^3^ | 1.05×10^3^ |  |
| Conjugation frequency (TC/D) | 9.21×10^-7^ | 9.17×10^-7^ | 2.50×10^-7^ | 6.96×10^-7^ |
