## Supplementary material for "GR13-type plasmids in *Acinetobacter* potentiate the accumulation and horizontal transfer of diverse accessory genes": Table S5

**Table S5. GR13 plasmids in GenBank (last search August 9, 2021)**

| **Plasmid** | **Accession** | **Host** | **Country** | **Year** | **Source** | **Size** | **L** | **+GR** | **antibiotic resistance genes** |
| --- | --- | --- | --- | --- | --- | --- | --- | --- | --- |
| p3ABAYE | CU459140 | *A. baumannii* | France | 2001 | human clinical | 94,413 | 1 | - | - |
| pMS32-1 | KJ616405 | *A. pittii* | China; Taiwan | - | - | 94,418 | 1 | - | - |
| p6411-89.111kb | CP010369 | *A. nosocomialis* | Colombia | 2012 | - | 89,111 | 1 | - | - |
| unnamed2 | CP014653 | *Acinetobacter* sp. | China; Panjin | 2015 | marine sediment | 50,047 | 1 | - | - |
| p2012N21-164-1 | CP033536 | *A. pittii* | China; Taiwan | 2012 | human | 97,329 | 1 | - | - |
| pC54_002 | CP042366 | *A. pittii* | Australia | 2014 | human clinical | 76,008 | 1 | - | - |
| p1_100020 | CP027251 | *A. pittii* | China; Chengdu | 2015 | human | 77,340 | 1 | - | - |
| p1_100004 | CP027247 | *A. pittii* | China; Chengdu | 2015 | human | 66,765 | 1 | - | - |
| pAP43-2 | CP043054 | *A. pittii* | China; Hangzhou | 2018 | human urine | 92,276 | 1 | - | - |
| unnamed3 | CP069540 | *A. pittii* | Germany | ≤1994 | human urine | 94,387 | 1 | - | - |
| unnamed2 | CP069506 | *A. pittii* | unknown | ≤1994 | human sputum | 94,379 | 1 | - | - |
| pAS61-1 | CP061590 | *A. seifertii* | China; Taiwan | 2010 | human blood | 93,205 | 1 | - | - |
| p2014N21-145-2 | CP033570 | *A. pittii* | China; Taiwan | 2014 | human | 72,034 | 1 | - | - |
| p2014S07-126-2 | CP033532 | *A. pittii* | China; Taiwan | 2014 | human | 96,775 | 1 | - | - |
| unnamed1 | CP077304 | *A. pittii* | Germany | ≤1994 | human ear discharge | 94,386 | 1 | - | - |
| p1_005069 | CP026087 | *A. pittii* | China; Chengdu | - | human | 91,563 | 1 | - | - |
| pDETAB5 | CP072528 | *A. baumannii* | China; Hangzhou | 2019 | intensive care unit environment | 97,035 | 2 | - | *bla*_NDM-1_, *ble*_MBL_, *bla*_OXA-58_, *aacC2d*, *msr*(E)*-mph*(E), *sul2* |
| pM131-2 | JX101647 | *A. soli* | China; Taiwan | 2010 | human sputum | 84,995 | 2 | *-* | *bla*_OXA-58_, *aphA1*, *aacC2d*, *sul2* (x2) |
| p255n_1 | KT852971 | *A. baumannii* | Vietnam | 2005 | human nasal | 92,939 | 2 | - | *bla*_OXA-58_, *aacC2d*, *bla*_VEB_, *arr-2*, *aadA1*, *aadB*, *cmlA6, msr*(E)*-mph*(E), *sul1*, *sul2* |
| pNDM1_010045 | CP028560 | *Acinetobacter* sp. | China; Chengdu | 2015 | sewage | 190,170 | 2 | 34 | *bla*_NDM-1_, *ble*_MBL_, *bla*_OXA-58_ (x2), *aacC2d*, *floR*, *msr*(E)*-mph*(E), *sul2*, *merA-2* |
| pOXA58_005078 | CP027245 | *A. baumannii* | China; Chengdu | - | human | 70,509 | 2 | - | *bla*_OXA-58_, *aacC2d*, *floR*, *msr*(E)*-mph*(E) |
| pBspH2 | CP055284 | *Acinetobacter* sp. | USA | 1986 | soil | 161,809 | 3 | - | - |
| unnamed3 | CP068198 | *A. johnsonii* | The Netherlands | 2008 | spacecraft-associated clean room | 206,659 | 3 | - | *-* |
| plas1 | CP070867 | *A. johnsonii* | China; Shanghai | 2018 | bigeye tuna | 149,408 | 3 | - | *-* |
| pAJ_082-3 | CP079794 | *A. johnsonii* | Pakistan | 2016 | intensive care unit sink | 155,957 | 3 | - | *-* |
| unnamed1 | CP033769 | *A. baumannii* | USA | 2016 | human sputum | 97,783 | - | - | - |
| pTS134338 | CP042210 | *A. baumannii* | India | 2005 | soil | 134,338 | - | - | - |
| pOCUAc17-1 | AP024799 | *A. pittii* | Japan* | ≤2021 | human blood* | 69,156 | - | - | - |
| unnamed1 | CP077233 | *A. pittii* | Germany | ≤1990 | human wound swab | 69,574 | - | - | - |
| pEC_gr13 | CP038263 | *A. baumannii* | Czech Republic | 2018 | frozen turkey liver | 128,013 | - | *-* | - |
| pE47_002 | CP042558 | *A. baumannii* | Australia | 2013 | hospital environment | 59,744 | - | - | - |
| pDETAB13 | CP073061 | *A. baumannii* | China; Hangzhou | 2019 | human rectal | 91,083 | - | - | - |
| pXBB1-8 | CP010358 | *A. johnsonii* | China; Chengdu* | - | sewage* | 117,483 | - | - | - |
| pCUVET596 | CP041149 | *A. baumannii* | Thailand | 2017 | dog urine | 82,016 | - | - | - |
| p2012C01-137-2 | CP033559 | *A. nosocomialis* | China; Taiwan | 2012 | human | 72,978 | - | - | - |
| unnamed1 | CP068175 | *A. ursingii* | The Netherlands | 2003 | spacecraft-associated clean room | 120,510 | - | - | - |
| pAR3 | CP038025 | *A. radioresistans* | Chile | 2008 | soil | 80,103 | - | - | - |
| p2014S06-099-1 | CP033541 | *A. pittii* | China; Taiwan | 2014 | human | 125,715 | - | - | - |
| pIC001A | CP022299 | *A. johnsonii* | Japan | 2003 | Tokyo Bay water | 94,476 | - | 26 | - |
| pCAM180A | CP044357 | *A. baumannii* | Cambodia | 2016 | human oral | 92,034 | - | - | - |
| p2010S01-197-2 | CP033563 | *A. nosocomialis* | China; Taiwan | 2010 | human | 92,044 | - | *-* | *bla*_OXA-58_ (x2), *aacC2d*, *sul2* |
| unnamed2 | CP069498 | *A. pittii* | Germany | ≤1994 | human blood | 128,321 | - | 24 | *-* |
| pEH_Gr13 | CP038259 | *A. baumannii* | Czech Republic | 2018 | human tracheal | 135,229 | - | - | - |

* = uncertain, information derived from submitter information and expected publication titles included in GenBank entries

L = lineage defined here, plasmids assigned to lineages 1, 2 or 3 are shaded pink, blue or orange, respectively

+GR = plasmids contain an additional replicon of GR type #
