## Supplementary material for "GR13-type plasmids in *Acinetobacter* potentiate the accumulation and horizontal transfer of diverse accessory genes": Table S6

**Table S6: Characteristics of the examined *dif* modules.**

| module | size  (bp) | putative function or content | reference  plasmid |
| --- | --- | --- | --- |
| *oxa58* | 2,257 | carbapenem resistance | pDETAB5 |
| *sul2* | 5,051 | sulphonamide resistance | pDETAB5 |
| *msrE-mphE* | 2,950 | macrolide resistance | pDETAB5 |
| *aacC2d* | 9,468 | aminoglycoside resistance | pDETAB5 |
| *dif*-696a | 696 | toxin-antitoxin | pDETAB5 |
| *dif*-696b | 696 | toxin-antitoxin | pDETAB13 |
| dif-823 | 823 | toxin-antitoxin (AdkAB) | pDETAB5 |
| dif-701 | 701 | toxin-antitoxin (HigAB) | pDETAB5 |
| *dif-*1491 | 1,491 | toxin-antitoxin + unknown ORFs | pDETAB13 |
| *dif*-2111 | 2,111 | alcohol tolerance | pDETAB13 |
| *dif*-6874 | 6,874 | alcohol tolerance | AP024799 |
| *dif*-7136 | 7,136 | alcohol tolerance | CU459140 |
| *dif*-28327 | 28,327 | copper resistance | CP068175 |
| *dif-*RND | 9,903 | RND efflux | pDETAB5 |
| *dif*-7932 | 7,932 | metabolism; possibly aromatic compound degradation | CP022299 |
| *dif-*1769 | 1,769 | sulphate permease | CP022299 |
| *dif*-6620 | 6,620 | small plasmid + mobilisation determinant | CP022299 |
| *dif*-801 | 801 | VRR-NUC domain protein | CP022299 |
| dif-614 | 614 | unknown | pDETAB5 |
| dif-1209 | 1,209 | unknown | pDETAB5 |
| *dif-*1612 | 1,612 | unknown | pDETAB13 |
